# Standardized Effect Sizes and Image-Based Meta-Analytical Approaches for fMRI Data

**DOI:** 10.1101/865881

**Authors:** Han Bossier, Thomas E. Nichols, Beatrijs Moerkerke

## Abstract

Scientific progress is based on the ability to compare opposing theories and thereby develop consensus among existing hypotheses or create new ones. We argue that data aggregation (i.e. combine data across studies or research groups) for neuroscience is an important tool in this process. An important prerequisite is the ability to directly compare fMRI results over studies. In this paper, we discuss how an observed effect size in an fMRI data-analysis can be transformed into a standardized effect size. We demonstrate how these enable direct comparison and data aggregation over studies. Furthermore, we also discuss the influence of key parameters in the design of an fMRI experiment (such as number of scans and the sample size) on (statistical) properties of standardized effect sizes. In the second part of the paper, we give an overview of two approaches to aggregate fMRI results over studies. The first corresponds to extending the two-level general linear model approach as is typically used in individual fMRI studies with a third level. This requires the parameter estimates corresponding to the group models from each study together with estimated variances and meta-data. Unfortunately, there is a risk of running into unit mismatches when the primary studies use different scales to measure the BOLD response. To circumvent, it is possible to aggregate (unitless) standardized effect sizes which can be derived from summary statistics. We discuss a general model to aggregate these and different approaches to deal with between-study heterogeneity. Furthermore, we hope to further promote the usage of standardized effect sizes in fMRI research.

## 1 Introduction

The analysis of neuroimaging data such as those obtained by functional magnetic resonance imaging (fMRI) is a complex endeavour. There are several challenges that need to be addressed. First there is a high cost to scan subjects. Second the data are typical of a high dimensionality and are characterized by a low signal to noise ratio (Button et al., 2013). Moreover, due to its complexity there is a high flexibility in the analysis and reporting of results (Carp, 2012). Given these challenges, it has been pointed out in literature that statistical power to detect common effect sizes in the neuroscience literature is rather low (Poldrack et al., 2017), resulting in an increased prevalence of false positive published effects (Ioannidis, 2005) and results that cannot be replicated (Patil et al., 2016) using an independent sample of new data (Turner et al., 2018). Several initiatives to address these issues have been proposed in literature. The first is the development of open source tools and standardization protocols to facilitate good research practices (Poldrack and Poline, 2015). For instance NeuroPower from the NeuroPowerTools^1^ can be used to calculate a minimal sample size to obtain sufficient statistical power when planning whole brain fMRI analyses (Durnez et al., 2016). Next the design of an fMRI experiment can be optimized (in terms of statistical efficiency) using NeuroDesign from the same software library. While these tools focus on planning an experiment, there is also an increased effort to standardize the organization and storage of obtained fMRI data. Examples are the Brain Imaging Data Structure (Gorgolewski et al., 2016) or the Neuroimaging Data Model (Maumet et al., 2016) which facilitates representation of meta-data. Combined, these protocols enable easy sharing of data between research groups.

A second solution directly addresses insufficient power by collecting data of thousands of individuals through large scale collaborations. Examples are the Human Connectome Project (Van Essen et al., 2013) or the UK Biobank (Sudlow et al., 2015). Finally, as an individual researcher it is possible to increase a sample size by aggregating data collected earlier (either published or not). Scientifically, this can be done using different meta-analytical techniques (Wager et al., 2007; Costafreda, 2009). The general idea is to synthesize results while weighting the different studies by reliability (e.g. studies with a larger sample size typically have a higher weight in a meta-analysis). Note that such initiatives are only possible when researchers engage in a data sharing culture (Poline et al., 2012).

In common between these solutions is the need to compare results over fMRI studies. This is not only an obvious prerequisite in order to combine results in a meta-analysis (Bowman, 2012; Borenstein et al., 2009) but also to gain insight into the magnitude of an effect. Most studies focus on null hypothesis significance testing, where the null hypothesis corresponds to the absence of an effect (Friston et al., 1995), and only report on the statistical significance. In case of fMRI data, an effect reflects the amplitude of the blood-oxygenated-level dependent (BOLD) signal obtained by contrasting (several) experimental conditions. The strength of the evidence in favor of the alternative hypothesis depends on the sample size as precision increases monotonically with the latter. Mathematically, it is true that any real effect (small or large) can be declared statistically significant as long as the sample size is large enough (Friston, 2012; Bzdok and Yeo, 2017). For this reason, the appropriate question is not only whether there is a true effect (and the direction of the effect) but also what the magnitude of the underlying effect is. This is termed practical significance (Kirk, 1996) and can be quantified by a (voxelwise) point estimate for the effect as well as an estimate of its precision. If effects are measured on a meaningful scale, then there is no need to standardize these (Cummings, 2011) before comparing or when combining over studies. However, the BOLD signal has an arbitrary unit (Chen et al., 2017) which creates a need to standardize fMRI effect sizes.

To our knowledge, the description of standardized effect sizes and an overview of methods to combine these estimates have not yet been fully documented in the field of neuroimaging. The goals of this paper are (1) to describe the advantages of standardized effect sizes, (2) to give an overview of the calculations and statistical properties of standardized effect sizes specifically related to fMRI data and (3) to discuss several methods to pool data across studies while dealing with heterogeneity between studies.

Note that several papers in literature describe various methods and tools to aggregate neuroimaging data (see for an overview: Wager et al., 2007) or provide specific guidelines on how to perform a meta-analysis with neuroimaging data (Müller et al., 2017). Most of these methods and diagnostic tools focus on coordinate-based meta-analyses that combine results over studies using only the 3D coordinates of peak activation from the individual studies (Acar et al., 2018; Albajes-Eizagirre and Radua, 2018; Eickhoff et al., 2009, 2012; Turkeltaub et al., 2002, 2012; Radua et al., 2012; Tench et al., 2017; Wager et al., 2009, 2004). In this paper, we focus on the context where full brain images are available and hence where information for each voxel is combined over studies. This is termed image-based meta-analysis. For a detailed comparison between both, see: Salimi-Khorshidi et al. (2009).

## 2 Effect sizes

We start this section with a more detailed argumentation to incorporate standardizing effect size estimates for fMRI data. We will contrast these estimates with the BOLD percent signal change. Thereafter, we will discuss the computations underlying standardized effect sizes and third, describe its relationship with fMRI data.

### 2.1 Why standardize?

The BOLD percent signal change is a frequently used measure of the magnitude of an effect (Chen et al., 2017). It is measured in response to an experimental condition and corresponds to the peak amplitude of the fMRI time series in relation to a baseline signal. The latter represents an average signal within each voxel over all time points. However, an average fMRI signal can vary between subjects or scanning sessions due to various external factors (e.g. caffeine level of the subject). Therefore, a normalization step is executed where the entire signal in each voxel within each subject and session is scaled to a target value. Written otherwise, if we denote *Y* as the observed signal, then the normalization corresponds to: 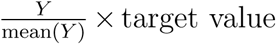. The target value ensures different sessions and/or participants are comparable *within* an analysis. The main issue is that there are incompatibilities between various software libraries for the analysis of fMRI data. More specifically, there are differences in 1) techniques to calculate an average signal (i.e. mean(*Y*)) and 2) the target values chosen to scale the signal. AFNI (Cox, 1996) for instance calculates the average signal within each voxel separately while SPM^2^(Statistical Parametric Mapping: Wellcome Department of Cognitive Neurology, London, UK) and FSL^3^ (Smith et al., 2004) perform a grand mean scaling where the average signal is calculated over all masked voxels (Chen et al., 2017).

With respect to scaling, SPM for instance sets the target value to approximately 100 while FSL (Smith et al., 2004) scales to ±10.000 due to historical differences in the implementation of the software^4^. Given that the BOLD percent signal change is calculated with respect to the baseline, the value depends on the technique to calculate the latter (i.e. software dependent) prohibiting direct comparison across studies using different software. In other words, consider two related studies that report a 3% BOLD signal change within an area of activation. The researchers of the first study relied on AFNI while the others on FSL. Unfortunately, the reported effects will be slightly different due to differences in the calculation of the average signal. Therefore, the BOLD percent signal change should be regarded as a non-standardized effect size. One of the solutions is to report standardized effect sizes (that are comparable across studies) next to the BOLD percent signal change.

To continue, we discuss the calculation of these standardized effect sizes and its relation to fMRI in section 2.3.

### 2.2 Standardized effect sizes

While standardized effect size calculations for a mean difference between two populations are well described (Borenstein et al., 2009; Zakzanis, 2001), this is not the case for a one sample average effect. In a group analysis of an fMRI study however, this is often of interest when pooling results over subjects. Therefore, we provide a general description for this specific effect size measure. Assume that the observed response variable *Y* for a given study *i* with *i* = 1, …, *k* is distributed as 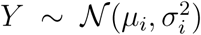. Here, *µ*_*i*_ is the true population effect of study *i* and 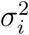 the corresponding within-study variance. The true standardized mean effect, *δ*_*i*_, is defined as:

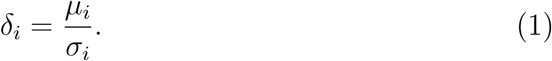

For ease of notation, we will drop the subscript *i*. Usually, *δ* is estimated using Glass’s estimator (Glass, 1976) or similarly for a one-sample case using Cohen’s *d* (Cohen, 1988):

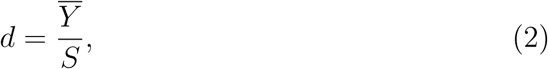

where *S* corresponds to the square root of the unbiased estimator for the sampling variability. Using equation (2) results in an expression of the observed effect 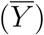 in units of *S*, which is a standardization procedure. However as shown in Hedges (1981) and demonstrated in section (5.1) of the Appendix, equation (2) is a biased estimator for *δ*. Relying on *d* results in an overestimation of *δ*, especially for small sample sizes. Using the derivations in Hedges (1981), the exact unbiased estimator (*g*_*e*_) is defined as:

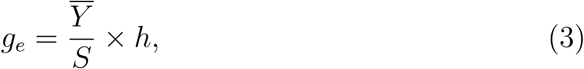

where

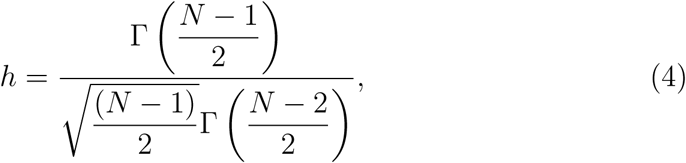

Γ represents the gamma function and *N* equals the sample size of the study. Due to historical limitations in computing *h*, Hedges (1981) suggested an approximation (*g*_*a*_):

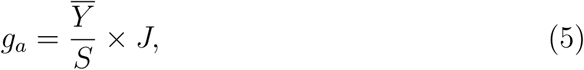

where

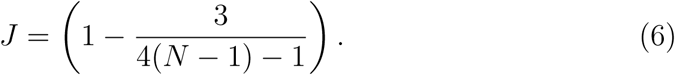

Note that for *N* ≥ 10, we have *g*_*a*_ − *g*_*e*_ < 0.001. Even though the difference between both effect size calculations is small, computational resources nowadays enable calculating *g*_*e*_ instead of *g*_*a*_. The induced overestimation using Cohen’s *d* for small sample sizes and the difference between *g*_*e*_ and *g*_*a*_ is illustrated in Figure 1. On the y-axis we plot the ratio between the true effect size (i.e. *δ*) over the expected value of *d* (i.e. E(*d*)) hence demonstrating the overestimation.

**Figure 1:**
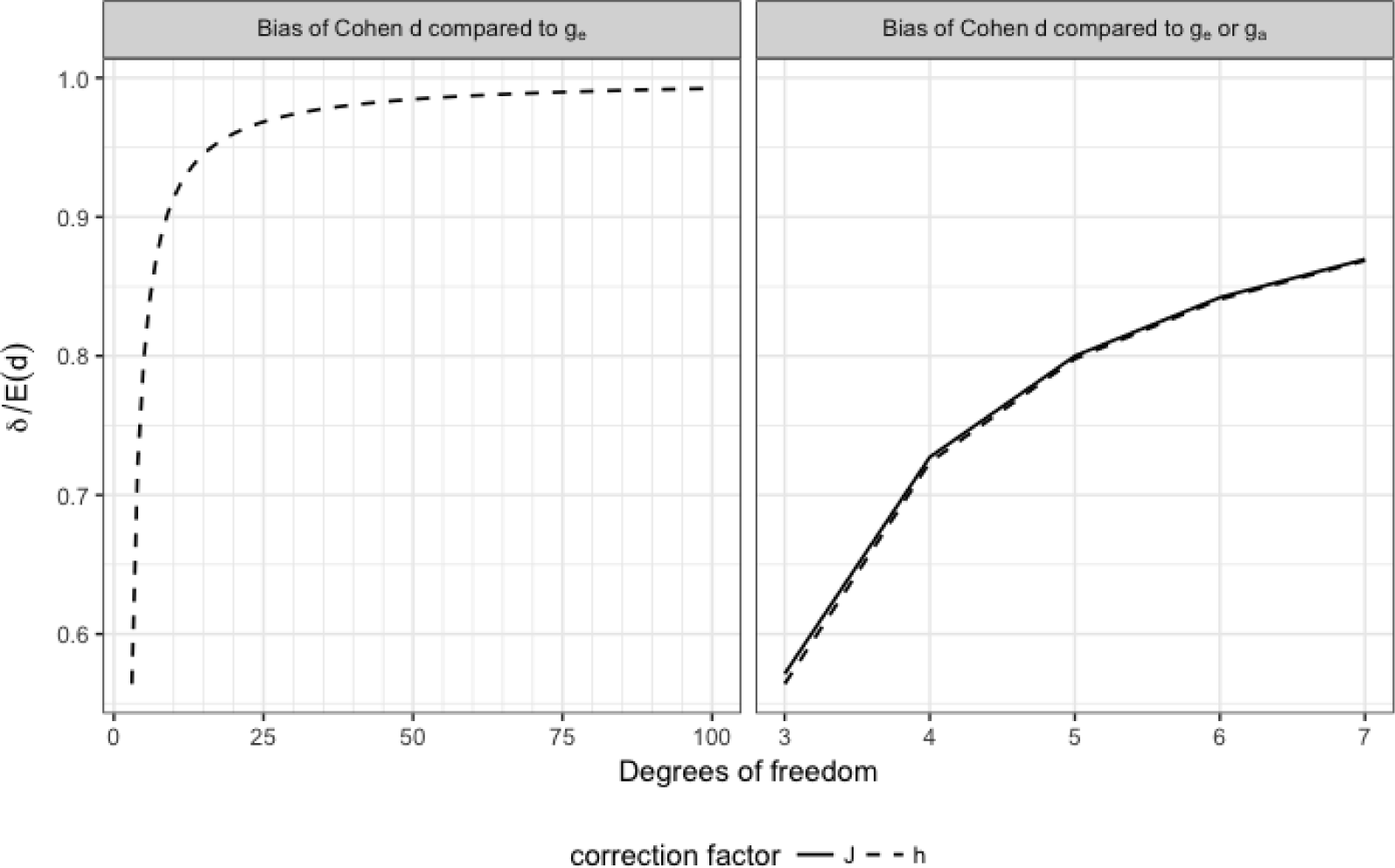
Left panel: ratio of the true effect size (*δ*) versus the expected value of Cohen’s *d*. Estimating an effect size with *d* thus results in an overestimation of *δ* for small sample sizes. The ratio equals the correction factor *h* used in the exact unbiased estimator (i.e. *g*_*e*_). The right panel shows the difference between the exact expression and the approximation (i.e. *g*_*a*_ using correction factor *J*) when the *df* ≤ 7. To summarize: 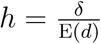 and 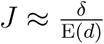.

Finally the expectation of *g*_*e*_ and its variance are given as:

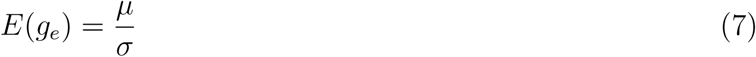

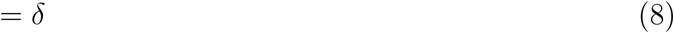

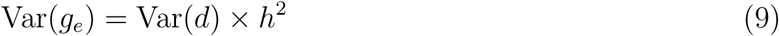

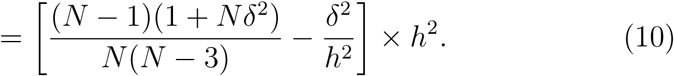

To estimate the within-study variability of the standardized effect, we plug in the estimate for *δ* using equation (2). We refer to the Appendix, section 5.1 for derivations.

For completeness, we provide the estimator for a standardized mean difference (*g*_Δ_) between a control group with *N*_*c*_ subjects and an experimental group with *N*_*e*_ subjects together with its expectation and variance (Hedges, 1984; Borenstein et al., 2009).

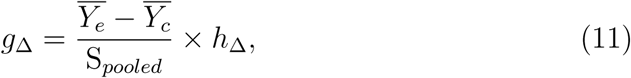

where

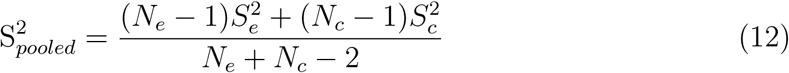

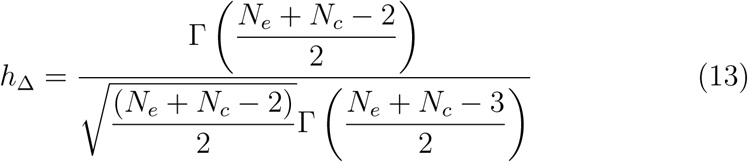

And then:

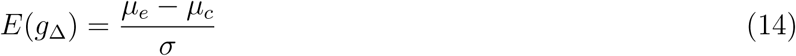

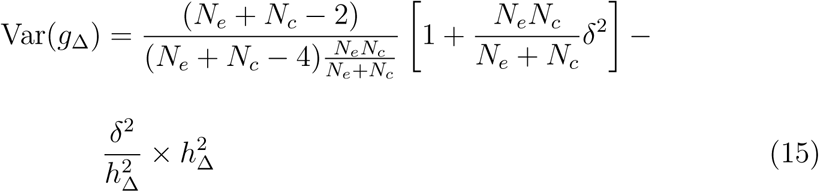

### 2.3 Standardized effect sizes for fMRI data

To discuss the relation between *g*_*e*_ and fMRI data, we need to give a brief overview of the general model fitting procedure that is typically used in a mass univariate fMRI data analysis.

#### 2.3.1 Statistical analysis of fMRI data

Due to computational limitations, the statistical analysis of fMRI data is split-up in two stages/levels. The first stage consists of fitting a general linear model (GLM) with the expected time series under the experimental condition and possible nuisance parameters to the observed BOLD signal. This is done for each voxel and subject separately. We will not use a subscript to denote the voxel for ease of notation. For a single subject *s*, we have (Friston et al., 1994):

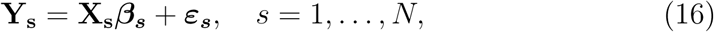

where **Y**_**s**_ is a vector of length *T* containing the BOLD signal on *T* measurements, **X**_**s**_ is the design matrix containing a convolution of the stimulus onset function with a hemodynamic response function (HRF) (Henson and Friston, 2007) as well as nuisance covariates, ***β***_***s***_ is a vector with the parameters and ***ε***_***s***_ is a vector of length *T* containing within-subject random error. Note that due to low-frequency drift, the within-subject error is not independently distributed but given by 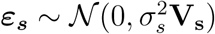, where **V** is the temporal autocorrelation (Mumford and Nichols, 2006). To obtain efficient estimators, we assume that the data, model and errors are decorrelated (whitened). This is usually done by premultiplying **Y**_**s**_ and **X**_**s**_ with a matrix **K**_**s**_ such that 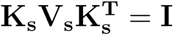 (Poline and Brett, 2012). By doing so, we have 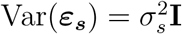 for each subject. Furthermore, note that for a fixed true ***β***_***s***_, we have

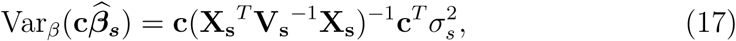

with 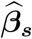 the vector of parameter estimators and where **c** represents a contrast vector forming a linear combination of parameter estimates (Mumford and Nichols, 2009).

The second stage, denoted with subscript *G*, is used to estimate a population effect by combining all participants. The response variable is the vector of estimated first level contrasts 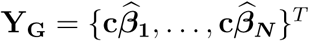. We now get

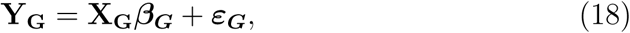

where **X**_**G**_ is a group design matrix and ***ε***_***G***_ a group error vector. In the simplest case, **X**_**G**_ corresponds to the group average which is a vector of length *N* with ones. Importantly, ***ε***_***G***_ is a mixed effects error component as it contains both variability of the imperfect intra-subject fit at the first stage (i.e. 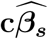) and between-subject variability, denoted as 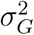 (Mumford and Nichols, 2006, 2009). To simplify the notation here, we look at the unusual case where each participant has the same design matrix **X** and we assume there is no temporal autocorrelation in the time series of each subject (or we work with pre-whitened data). We thus have 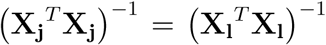 for all *j, l* after which we drop the subscript in **X**. The variance of the error component in model (18) becomes:

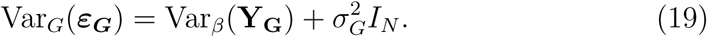

with

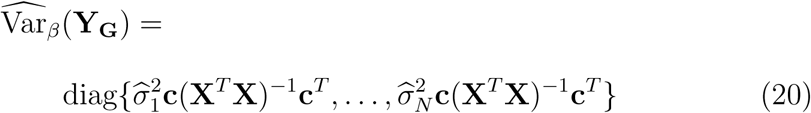

where diag indicates the diagonal operator and *β* denoting this is an intra-subject variance.

Next we assume that the within-subject variance is homogeneous over all subjects (i.e. 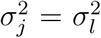 for all *j, l*). This simplifies our discussion on standardized effect sizes in the following section. Furthermore, by doing so we are able to rely on the ordinary least squares (OLS) approach (Holmes and Friston, 1998) to estimate the parameters of equation (18). Finally, we get for a fixed true ***β***_***G***_:

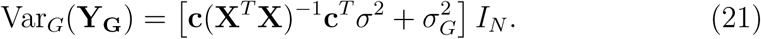

We only need to estimate a single (combined within- and between-subject) variance term. We shall denote 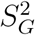 as the sample variance of *Y* at the group level (i.e. 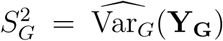). Inference for the simple group average case proceeds with a one sample *t*-test. The test statistic (i.e. a *t*-value) of a parameter of interest (i.e. a specific *β*_*G*_) corresponds to 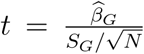 and is compared to a *t*-distribution with *N* − 1 degrees of freedom. The result is a statistical parametric map (SPM) containing a test statistic for each voxel.

#### 2.3.2 Model fitting procedure and effect size estimation

While it is possible to calculate a voxelwise standardized effect for each subject in an fMRI experiment, the general interest is to measure a standardized group effect. Specifically for fMRI, the one-sample standardized mean effect (i.e. **X**_**G**_ = a column vector of 1) is obtained by combining equations (3), (18) and (21) as:

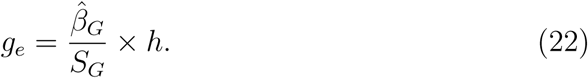

Note that it is also possible to obtain an effect size using the *t*-value in each voxel. More particularly, we have 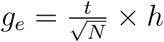.

Due to the two-stage procedure when fitting the GLM, there are some characteristics/effects on the standardized effect size worth noting. To start with, consider the expression of *δ* which contains both an estimate of the average effect as well as one of the sampling variability. Therefore, both estimators for *µ* and *σ* need to be unbiased. For instance, we make the assumption that the individual time series are effectively decorrelated. Furthermore we need to assume the correct statistical model is used to estimate the parameters of the GLM (e.g. movement parameters are included if necessary). In other words, we assume researchers are measuring a relevant effect instead of an artefact. Second, the derivations for the distributional properties of the standardized effect sizes are obtained using the assumption of a normally distributed outcome variable. In the case of fMRI, we thus assume normally distributed *β*_*G*_’s, which follows from the central limit theorem if *N* is sufficiently large.

Next, there is an effect of both the length of the individual time series (*T*) and the number of participants (*N*) on the standardized effect size and its variance. First note that the sampling variability 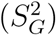 will decrease as *T* increases. This is the case as the intra-subject variability due to estimating the single-subject parameters in the fist stage (i.e. the first part in equation 21) decreases. In turn, *δ* will increase. Thus one obtains larger effect sizes using longer scanning sequences. However note that larger effect sizes are associated with larger variances of the effect size estimator as can be seen in equation (10). Secondly, while the effect size is independent of the sample size, the precision to estimate *δ* will increase with *N*. To demonstrate both the effect of *T* and *N*, we set up a small Monte-Carlo simulation study. The code for this simulation and to obtain the resulting figures can be found at: https://github.com/NeuroStat/ESfMRI. Using the *neuRosim* package (Welvaert et al., 2011) in `R`, we generate individual fMRI BOLD time series by convoluting a block design experiment with a canonical HRF and add white random noise to the signal. Within-subject variability is homogeneous across all subjects and a small amount of between-subject variability is added. We then vary the length of the individual time series with *T* ∈ [100, …, 500] and let *N* = 20 or *N* = 100. A standard two-stage GLM is fitted and we estimate the group level standardized effect size with its variance using the formula provided above. Results are shown in Figures 2 and 3. First we visualize standard properties of the OLS estimator for the parameters of interest in the second level GLM. The estimator for ***β***_***G***_ is unbiased and its precision increases with both *T* and *N* (i.e. lower variability over the simulations). Second we observe higher values for *g*_*e*_ as *T* increases and a higher precision as *N* increases. Furthermore, the variance of *g*_*e*_ increases with *T*. This can be seen in panel **D** of Figure 5.3 where the red line indicates the expected value for the variance. Note that the increase is rather small. Indeed, the empirical variance (i.e. over Monte-Carlo simulations) of the estimator for *g*_*e*_ increases, albeit this is too small to be detected in panel **C** of Figure 5.3. A demonstration of the increase in the empirical variance is given in Appendix, section 5.2, where we calculate the variance over the simulations of the estimates for *g*_*e*_ marginally over the number of subjects. Note that even though the empirical distribution of Var(*g*_*e*_) is skewed (see panel **D** of Figure 5.3), the estimator is unbiased. A demonstration of this property is given in the Appendix, section 5.3. Furthermore, note how the variance for *g*_*e*_ decreases with the number of subjects. This can be seen in both panel **C** and **D** of Figure 5.3 and is calculated explicitly in the Appendix, section 5.2. This effect is more substantial than the increase in the variance associated with the number of scans.

**Figure 2:**
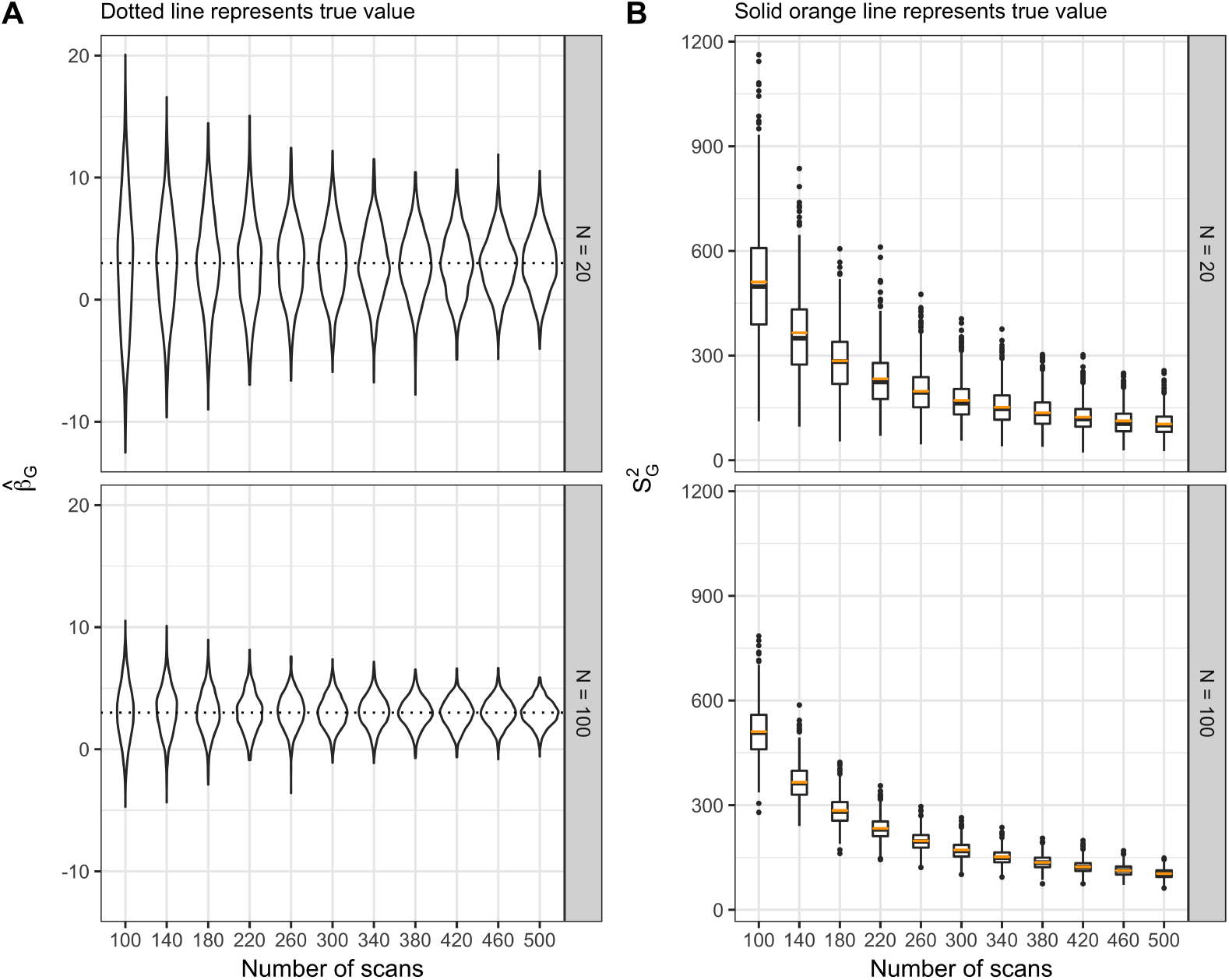
Empirical distributions over 1000 Monte-Carlo simulations of the estimated group level parameters in a typical two-stage fMRI general linear model (see equation (18)). The length of the individual time series and the total sample size increases to demonstrate its effect on the parameter estimates. Panel **A** depicts the estimated group level parameters, while panel **B** gives 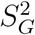.

**Figure 3:**
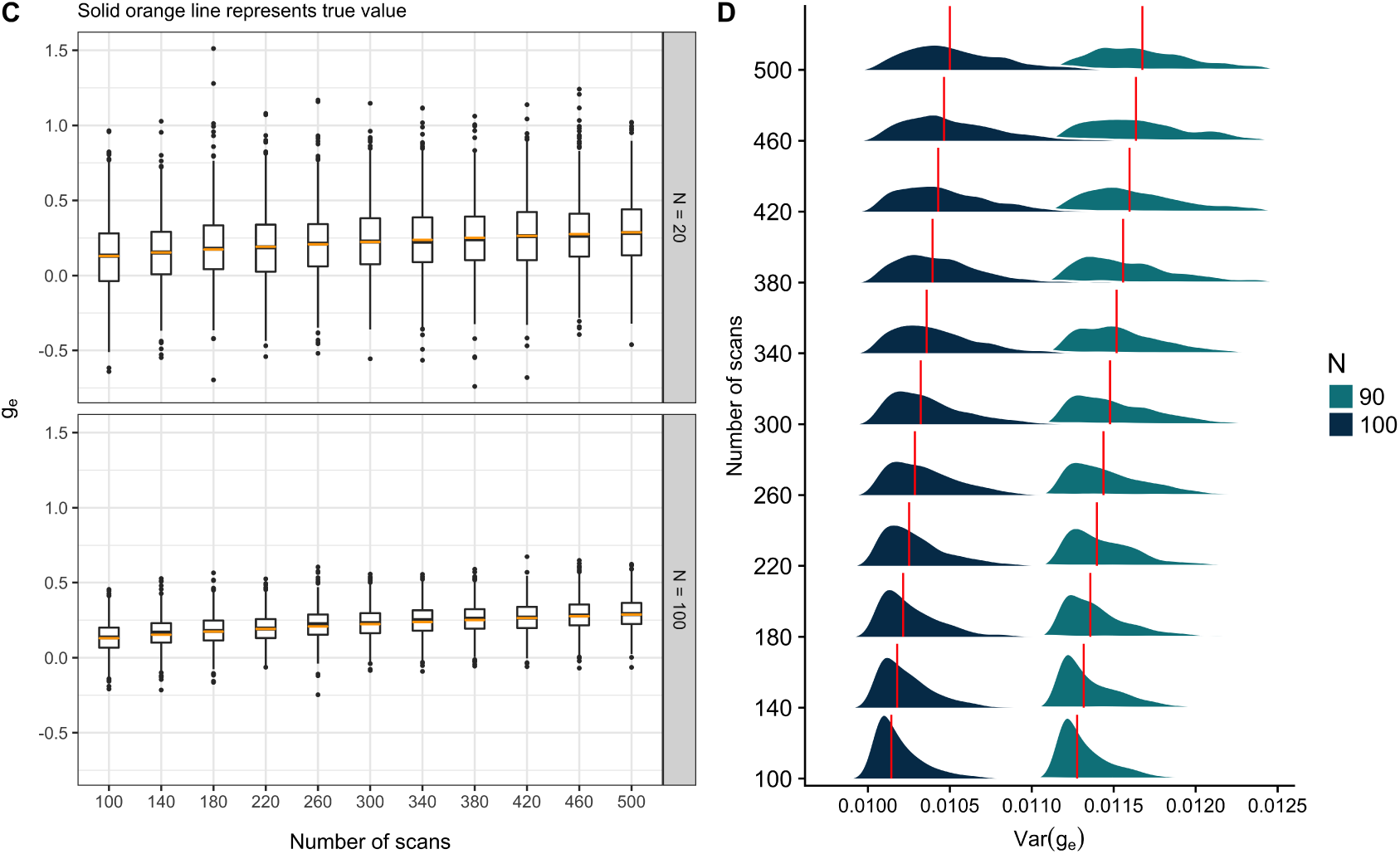
Empirical distributions over 1000 Monte-Carlo simulations of the estimated *g*_*e*_ (panel **C**) and its variance (panel **D**) using equation (10). The length of the individual time series and the total sample size increases to demonstrate its effect on the parameter estimates. The solid red line in panel **D** corresponds to the expected value.

Finally, there will be an effect of spatial smoothing in the pre-processing on the estimated standardized effect size. As smoothing directly influences the signal and noise of within-subject time series, the smoothed standardized effect size will be different from the raw standardized effect.

An important remark is the distinction between effect size estimation and null hypothesis significance testing for fMRI. Both procedures are mainly used within a massive univariate context (Lindquist, 2008). However, inference procedures for the parameters of the GLM critically relies on correcting for multiple testing (Nichols and Hayasaka, 2003; Nichols, 2012). While effect sizes can be calculated on thresholded maps, the resulting estimates will be over-optimistic as the amount of tests increases. This is true as only voxels with the smallest amount of noise will survive the threshold for significance, irrespective of the underlying true effect size (Reddan et al., 2017). Therefore, we suggest to complement significance testing with estimating the standardized effect sizes on non-thresholded images.

## 3 Meta-analysis

There are several methods available to aggregate fMRI data. However, the amount of information available to the analyst will largely determine which method can be used. We discuss two strategies aggregating results when full brain images are available (i.e. image-based meta-analysis). In an fMRI meta-analysis, three levels can be distinguished: subjects, studies/groups and the meta-analytical level.

In the first method, full brain images from different studies are modeled by extending the two-stage hierarchical general linear model presented in section (2.3.1). To do so, the analyst needs the parameter estimates of the GLM of each study (equation 18) together with an image containing the estimate of its variance. Furthermore, depending on the software used within each study, meta-data will be required to scale data in order to avoid unit mismatches between studies. The second method relies on a transformation of the estimates from each study to standardized effect sizes (discussed in section 2.3.2). This approach can be used if only images containing the test-statistic of each study are available.

### 3.1 Extending the GLM

We start with the case where most information is available. It is possible to extend the two-stage hierarchical GLM presented in equation (18) with a third level. To do so, we need at least the following information from each study *i* with *i* = 1, …, *k*: the estimated contrast parameters of interest from the GLM 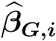 together with their estimated variances, the sample size or preferably the effective degrees of freedom for each voxel and finally meta-data such as the mean intensity of each study image. If we use *M* to denote the third (meta-analytical) level and **c**_**G**_ as a contrast forming vector at group level (e.g. contrasting patients with controls) in each study, then we have 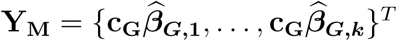. The corresponding GLM can be written as:

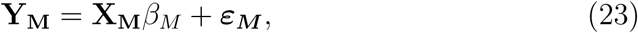

where **X**_**M**_ represents the meta-analytical design matrix used to aggregate data. Similar to the two-stage model presented earlier, the error term consists of two parts. In this case: within-study variability and between-study variability 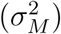. Note however that within-study variability on its own consists of the intra-subject and between-subject variability. Written formally, we thus have:

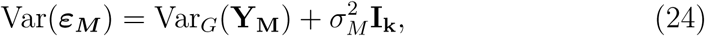

with

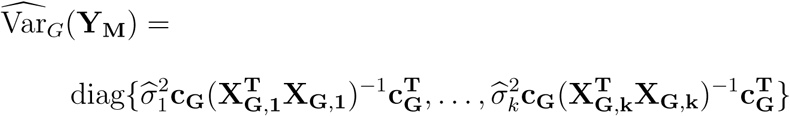

and diag again indicating the diagonal operator and for each study 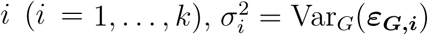 as in expression 19. Note that differences in sample sizes may cause within-study variability being non-homogeneous. In turn, this violates an assumption underlying the OLS approach to estimate the parameters of model 23. As explained in Mumford and Nichols (2009), it is possible to relax the assumption of homogeneous within-study variances by using a generalized least squares approach. Since this requires an estimation of multiple variance terms, we need to use iterative algorithms such as restricted maximum likelihood. For fMRI, it is possible to use a general linear mixed model implemented as *FLAME1* or *FLAME1*+*2* in FSL (Woolrich et al., 2004) or the approach implemented in fMRIstat (Worsley et al., 2002). Note that if the assumption of within-study homogeneity is violated, one should avoid the summary statistic approach in SPM12^5^. However, it is possible to use a full general linear mixed model implemented as *spm_mfx.m* in SPM12 and described in (Friston et al., 2005).

While modeling the subject- or study-level data using a hierarchical three-stage GLM seems straightforward, special attention needs to be given to the units of the data. Ideally, the effect of an experimental paradigm (i.e. the *β* parameters in the GLM) is expressed in the same unit for each study before going into model (23). However as mentioned in section 2.1, the BOLD signal has an arbitrary unit. This will not only depend on the scaling technique used when normalizing the time series (as discussed earlier), but also on the scale of the design matrix and the scaling of the contrast of interest at the first level of the GLM in each study (Maumet and Nichols, 2015). With respect to the design matrix, ideally predictors have a peak amplitude (height) of 1 ensuring the effect is expressed in the same unit as the data. This is usually the case for block event designs, but rarely for event related designs with short inter-trial intervals. With respect to the scaling of the contrast, ideally the positive part sums to 1 and the negative part sums to −1 (see also https://blog.nisox.org/2012/07/31/spm-plot-units/). In most cases, the analyst will not be able to change these parameters. Avoiding unit mismatches between studies will therefore become challenging.

To demonstrate this, assume that two individual fMRI studies rely on the same technique to calculate an average signal and both studies are interested in the same experimental contrast, measuring a relevant effect. As discussed above, a typical data analysis consists of fitting a general linear model (GLM) in each voxel. Even though both studies could measure the same BOLD percent signal change, the parameter estimates in the GLM will differ if the target value of the normalization step differs. To compare these parameter estimates over both studies (possibly with the aim to aggregate them in a meta-analysis), one then needs to use an extra parameter to scale the design matrix from one study to another. An illustration of this problem is given in Figure 4. Using `R` (R Core Team, 2015), we run a simulation^6^ where we generate 1000 time series for *N* = 50 subjects each. We design two scenarios. These correspond to the signal being normalized with a target value in the normalization step corresponding to either 10.000 (option A) or 100 (option B). We demonstrate how a difference in normalization results in parameter estimates that cannot be combined unless the analyst knows how to scale data between studies (using a scaling parameter ***c*** on Figure 4).

**Figure 4:**
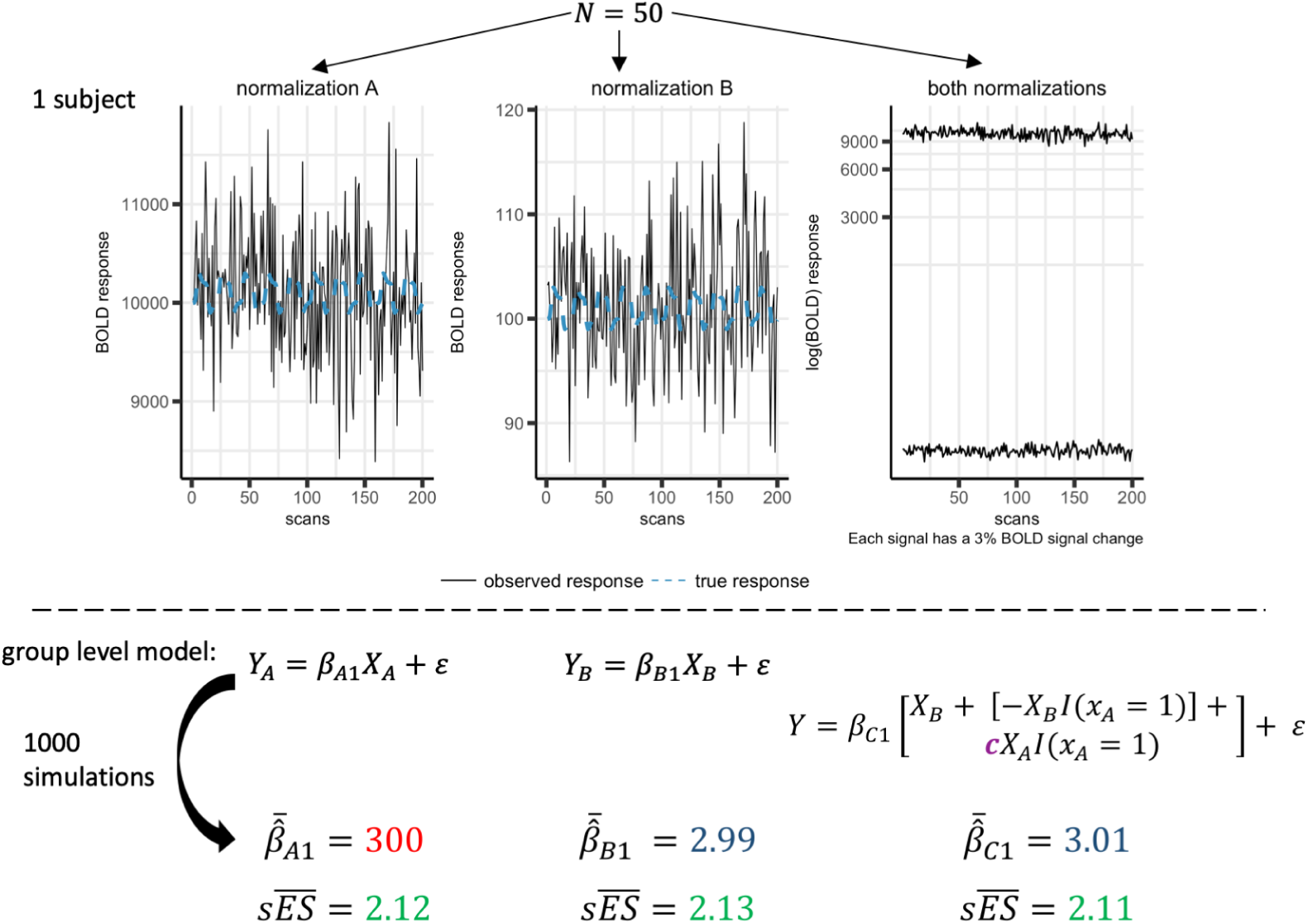
Illustration of pooling two studies (*A* and *B*) where each have a different target value in the normalization step. This results in regression coefficients (i.e. *β*_*A*1_ and *β*_*B*1_) that cannot be compared directly. Two solutions are possible: 1) one can scale the design matrix (i.e. *X*) of one study to the other (resulting in *β*_*C*1_) or 2) one could standardize the observed effect size (i.e. *sES*). To generate the observed time series, we first create a true signal by convoluting a block design experiment (20 sec ON/OFF) with a canonical hemodynamic response function (HRF) (Henson and Friston, 2007). The true signal is scaled so that it corresponds to 3% BOLD signal change relative to the baseline. We then add white random noise to the true signal and a small amount of homogeneous between-subject variability. Note: ***c*** corresponds to the scaling parameter needed to combine study *A* and *B*.

If raw data (i.e. subjects) are available, one could re-run the entire analysis within one software library thus avoiding differences in scaling techniques between software libraries and with the ability to re-code contrasts vectors. Furthermore, one could obtain for each study the effective peak height of the predictors in the design matrix and use this value to scale the parameter estimates accordingly before fitting the second-level GLM.

If only the second-level group data are available (i.e. parameter estimates and their estimated variances), the analyst will need to rely on meta-data (e.g. software library used, type of experiment, the chosen contrast of interest,…) to scale the estimates accordingly before running a third-level. Standardization protocols such as the Neuroimaging Data Model (Maumet et al., 2016) attempt to provide machine-readable descriptions of studies together with the necessary meta-data to overcome these issues.

If no meta-data is available or only a limited amount of information is available (e.g. only the statistical parametric map), then the analyst has to rely on techniques that do not depend on the unit of the data. We will discuss these now.

### 3.2 Aggregating standardized effect sizes

An alternative method to aggregate fMRI studies relies on transforming the data beforehand. As discussed in section 2.3.2, it is possible to transform a test-statistic to standardized effect sizes. Furthermore, note that the non-systematic variability of standardized effect sizes is proportional to the inverse of the sample size 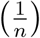 of the study (see equation 10). Moreover, the variance is completely determined by the estimate of the effect size and the sample size. This has three important implications. First, effect size estimates have a different error variance due to different sample sizes between studies (Hedges, 1984). These differences are accounted for when aggregating effect sizes by using a weighting approach (discussed below). Second, aggregating studies is more feasible as one only needs the estimate of the effect magnitude (obtained using a test statistic) and the corresponding study sample size. For fMRI it is thus possible to do a meta-analysis using the statistical parametric maps in conjunction with the sample sizes. Finally, it is possible to use all degrees of freedom between the different effect size estimates to estimate systematic differences between studies (Hedges, 1984). In other words: there is no need to iteratively estimate within-study and between-study variance components as could be the case in the three-stage GLM from equation (24).

#### 3.2.1 Models for aggregating effect sizes

In the past, it was custom to ignore any possible between-study variability (i.e. fixed-effects models) when aggregating effect sizes. However there is a growing consensus that more complex models are preferred over these simple fixed-effects models (Hunter and Schmidt, 2000; Viechtbauer, 2005). Similar to the three-stage hierarchical model, variability of effect sizes are then decomposed into two parts. The first is the sampling variance (i.e. within-study variability) determined by the sample size and the second equals heterogeneity between studies due to observed effect sizes being drawn from a population of effect sizes (i.e. between-study variability). Typically, models incorporating the latter are denoted as a random effects model. However we will discuss a more general case which allows for moderator variables to be added. We introduce new notation to differentiate models for standardized effects from those in BOLD percent signal change discussed earlier (section 3.1). We can write a meta-analysis for all *i* = 1, …, *k* studies as a linear mixed-effects model (e.g. Raudenbush, 2009; Viechtbauer et al., 2015):

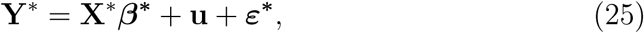

where **Y**\* denotes the (*k* × 1) vector of observed (standardized) effect sizes and **X**\* is the *k* × (*p* + 1) meta-analysis design matrix with *p* possible moderator variables. Next, ***β***\* is the column vector of length (*p* + 1) with parameters, **u** is the vector of random effects distributed as 𝒩 (0, *τ* ^2^). Then *ε*\* is the within-study error with the individual terms distributed as 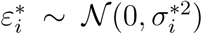. The mentioned random effects model without moderator variables corresponds to the case where **X**\* is a column vector of 1’s. The intercept in ***β***\* then corresponds to the population average. Furthermore *τ* ^2^ reflects all between-study heterogeneity not accounted for by the moderators. A weighted least squares approach is used to estimate the parameters in ***β***\*. The usual estimator for ***β***\* is denoted by **B**\* and is expressed as:

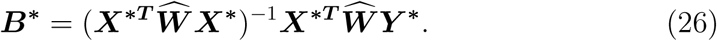

In this, 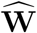 denotes a diagonal weight matrix with elements 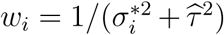. We thus give less weight to studies with a lower sample size (i.e. lower precision).

A great deal of attention in the literature deals with methods to estimate *τ* ^2^, the between-study heterogeneity (Higgins and Thompson, 2002; Veroniki et al., 2016; Viechtbauer, 2005). Choosing an appropriate method to estimate *τ* ^2^ is important for two reasons. First, a biased estimator leads to an over- or underestimation of ***β***\*. Second, the variance-covariance matrix of **B**\* used in a standard Wald-type test can be expressed as 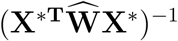 (Viechtbauer et al., 2015) and hence, the precision to estimate the parameters is also a function of *τ* ^2^.

In the following section, we will discuss three methods that are worth mentioning to estimate *τ* ^2^ for fMRI data. These are the popular DerSimonian and Laird estimator (DerSimonian and Laird, 1986), the Hedges estimator and the restricted maximum likelihood estimator. Moreover, we will discuss these using an empirical assessment of the amount of between-study heterogeneity that one can expect in an fMRI meta-analysis. Obtaining such an estimate has three benefits. First it can be used in subsequent Monte-Carlo simulation studies by providing sensible values for simulation parameters. Second, results of these assessments challenge the analyst to think ahead about potential between-study heterogeneity when planning a meta-analysis. Finally, the expression for *τ* ^2^ can attain negative values at which they are usually truncated at zero. This leads to biased estimates (Bohning, 2002). Hence it is interesting to know whether one can encounter this situation in an fMRI meta-analysis. We collect a real data set of 33 studies to discuss the estimators for *τ* ^2^ and the observed distribution of between-study heterogeneity within a region of interest.

#### 3.2.2 Between-study heterogeneity in fMRI studies

We collect a database of whole-brain fMRI studies involving a general experience of *pain* versus *no pain*. We obtain 33 studies using NeuroVault where we transform the SPMs to standardized effect sizes. We then estimate between-study heterogeneity using 3 possible estimators for *τ* ^2^ (discussed below). Furthermore we add two measurements that quantify the degree of between-study heterogeneity. We plot all the observed values over all voxels within a region of interest in Figure 5. The different steps are detailed below and the code is available at our GitHub repository.

**Figure 5:**
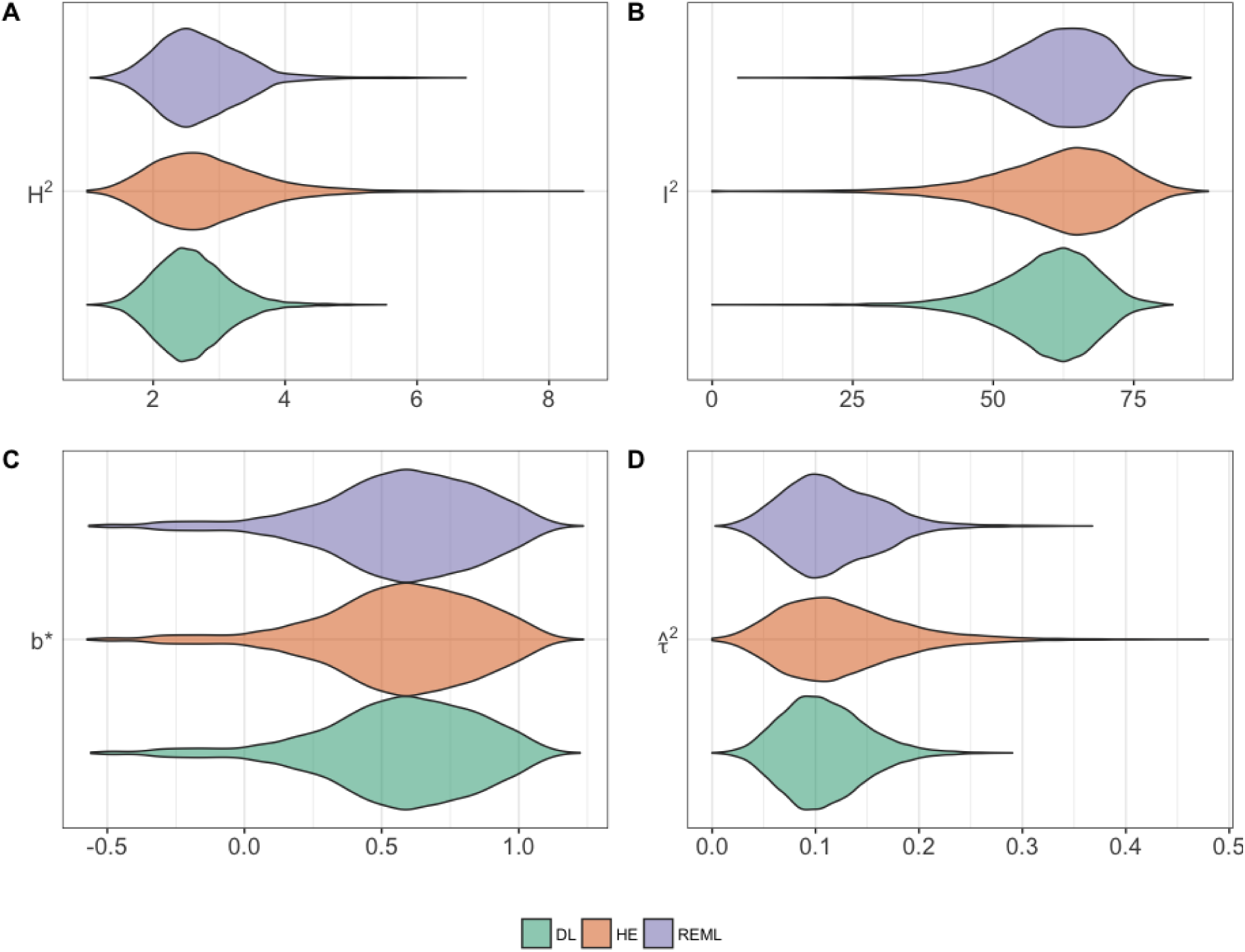
Empirical distributions for various parameters obtained after fitting a moderator meta-analysis for the contrast PAIN > NO PAIN within a ROI. Panel **A** and **B** visualize two measurements for the amount of between-study heterogeneity based on Hedges estimator. Panel **C** shows the estimated effect in standardized units. Panel **D** depicts the estimated between-study variability. To estimate the latter, we use the DerSimonian and Laird (LD) estimator, the Hedges (HE) estimator and an approach using the Restricted Maximum Likelihood (REML).

##### Step 1: collecting studies

All studies are collected using the search term *pain* in NeuroVault (last search dates from January 2017). The contrast of interest is the effect of experiencing pain versus a baseline or versus experiencing no pain. We apply a lenient definition of the experience of pain to obtain upper bounds on the observed between-study heterogeneity. The database contains multiple stimuli, ranging from auditory, thermal to mechanical stimuli. Results are manually checked for content and/or errors. If needed, images are re-sampled to MNI space with a voxel size of 2 × 2 × 2 mm. Note that we do not explicitly follow guidelines (such as the PRISMA statement^7^) on how to collect/report studies for a meta-analysis/systematic review. Nor do we aim to assess whether publication bias known as the ‘file drawer problem’ (Rosenthal, 1979) is present. Hence the research question is not necessarily on the mapping of the experience of pain in the human brain. The database contains 33 studies from which 21 correspond to the work of one research group. Furthermore, 6 studies provide individual subject data containing only the beta estimates of the fitted GLM. We re-create the studies for these by fitting group level models using a standard ordinary least squares approach (Holmes and Friston, 1998), assuming homogeneous within-subject variability. The total sample size equals 659 subjects.

##### Step 2: defining a region of interest

As is done in Poldrack et al. (2017), we restrict our analysis to regions that are functionally relevant to the experience of pain. To obtain a region of interest (ROI), we run an automated meta-analysis using NeuroSynth^8^ (Yarkoni et al., 2011). This tool automatically extracts the coordinates of foci of activation from published papers. Out of the entire database of published fMRI studies, 420 studies reported activation associated with *pain*. For each voxel, a cross table *mentioned pain in a study (YES/NO)* × *reported activation (YES/NO)* can then be constructed. Next the probability that activation in a given brain voxel is associated with the term *pain* is calculated (i.e. P(activation|*pain*)). The Pearson *χ*^2^ test is used to test the null hypothesis of independence between each brain voxel and *pain*. A binary mask (i.e. functionally relevant: 1/0) is then created by controlling the false discovery rate at level 0.01.

##### Step 3: estimating between-study heterogeneity

Using the *NeuRRoStat* package^9^ in `R`, we first transform the *t*-values within the ROI to standardized effect sizes (i.e. *g*_*e*_) based on equation (3). Within-study variability is calculated using equation (10). Next we use the *metafor* package (Viechtbauer, 2010) in `R` to fit a mixed-effects model as given in equation (25) where a fixed moderator variable is added to control for the 21 studies corresponding to the same research group. The weighted least squares approach requires an estimation of *τ* ^2^ - the between-study variability. To condense the amount of formula in the text, we refer to the appendix section 5.4 for the mathematical description of the three estimators for *τ* ^2^.

The first is the popular DerSimonian and Laird (DL) estimator (DerSimonian and Laird, 1986). This method of moments estimator has the advantage of being very quick to calculate. However, as discussed in Viechtbauer (2005), the estimator is only unbiased assuming that within-study variances (*σ*^*2^) are known (i.e. not estimated). Obviously this assumption does not hold and becomes especially problematic if the sample sizes of individual studies are small. Furthermore, even though the estimator does not require an assumption of normality for the random effects, it has been shown in a simulation study how other methods are preferred when the amount of studies in the meta-analysis is small (Jackson et al., 2010). A small adaptation of the DL estimator is given by the Hedges estimator (HE) (Viechtbauer, 2005). This estimator is proven to be unbiased when assuming the within-study variances are estimated rather than being known and has the same computational advantage as the DL estimator. However as shown in Viechtbauer (2005), the HE estimator is the least efficient, meaning the variance of the estimates is the highest among the three estimators. A final alternative is an iterative approach using the restricted maximum likelihood (REML). This estimator benefits from a good compromise between efficiency and being unbiased. The disadvantage is the increased computational time and possible convergence problems. We estimate *τ* ^2^ using all three approaches presented above.

##### Step 4: visualizing between-study heterogeneity

To complement the analysis, we add two commonly used estimates of between-study heterogeneity. The first one being *H*^2^ which is defined as 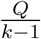, where *Q* is the amount of observed dispersion between studies. Note that *Q* on itself depends on the estimator for *τ* ^2^ and *H*^2^ should therefore not be considered as an independent check of heterogeneity. The expected value of dispersion under the assumption of homogeneous effects (i.e. there is no between-study heterogeneity) equals *df* = *k* −1 (i.e. equal to the degrees of freedom). Therefore, *H*^2^ is a measure of the ratio of observed dispersion over the expected amount of dispersion. We also add *I*^2^ which is defined as 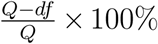 and thus the ratio between excess dispersion over the total (observed) dispersion. We visualize the distributions for these various parameters within the ROI in Figure 5. We observe values for *H*^2^ and *I*^2^ that indicate substantial between-study heterogeneity. For instance, the maximum values for *H*^2^ and *I*^2^ using the Hedges estimators equal 8.52 and 88.3% respectively. There is no rule of thumb to describe the observed heterogeneity as either low, mild or severe (Higgins and Thompson, 2002). However, we believe the observed dispersion can be considered substantial. With respect to the different estimators for *τ* ^2^ (i.e. DL, HE and REML), we note the following observations. First, the HE estimator is associated with more outlying estimates of *τ* ^2^, followed by the REML estimator. However, this does not seem to affect the estimation of ***β***^*^ as these distributions are nearly identical. The range of *B*^*^ over the three estimators is between −0.573 and 1.24 (panel **C** of Figure 5). Note that *τ* ^2^ is estimated on the same scale. Furthermore, we can construct a 95% credibility interval of the estimated *B*^*^’s (for each estimator of *τ* ^2^) using the assumption of normally distributed effects around *B*^*^ (i.e. indicating the amount of variability one can expect). For instance, using the HE estimator, the interval associated with the voxel with the largest estimated *B*^*^ (i.e. 1.24) in the ROI is equal to [0.385; 2.09]. The interval for the lowest observed *B*^*^ (i.e. −0.573) equals [−1.36; 0.218]. As a reference, the interval for the voxel with the highest value for 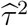 (i.e. 0.48) using HE equals [−0.891; 1.82]. Written otherwise, there is at least some between-study heterogeneity present that cannot be explained by the moderator variables included in the analysis. Moreover, we observe only a few number of zero-valued estimates. This implies we do not need to worry about bias induced to truncating the variance estimates to zero. Finally we plot the observed effects together with the estimates for 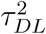 on an 2 × 2 × 2 MNI template in Figure (6). We can see how larger values for *B*^*^ are associated with larger values for 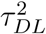.

**Figure 6:**
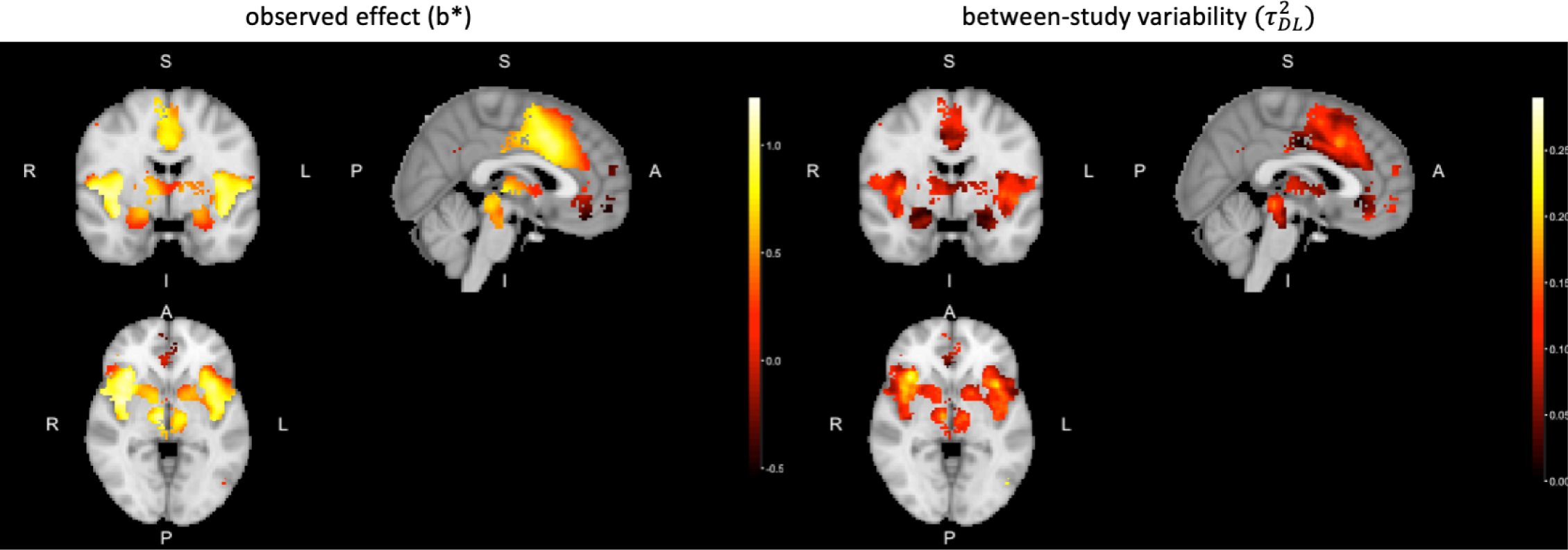
Estimated weighted effect (*B*^*^) for the PAIN > NO PAIN meta-analysis on the left and corresponding between-study variability (*τ* ^2^) using the DerSimonian and Laird estimator on the right for voxels within the ROI. The MNI coordinates of the slices correspond to *x* = 0, *y* = 0 and *z* = 0. Larger estimates for *B*^*^ are associated with higher between-study heterogeneity.

To conclude, we show how a typical fMRI meta-analysis will need to deal with a substantial amount of between-study heterogeneity.

#### 3.2.3 Alternative methods

While transforming a test statistic to standardized effect sizes before aggregating is one way to avoid differences in the unit between studies, alternative methods are possible. These focus on combining test statistics as these have no unit either. For instance, Fisher’s combined probability method (Fisher, 1925) converts *Z*-values from independent tests (i.e. the different studies) to *P* -values and then combines these into a *χ*^2^ test-statistic. Another approach is Stouffer’s method and can be used to average *Z*-statistics while also weight studies by their sample size (Liptak, 1958). Furthermore, it is also possible to avoid distributional assumptions when performing statistical inference using non-parametric methods such as permutation testing. An overview of these methods together with simulation results tailored to fMRI data are given in (Maumet and Nichols, 2016).

## 4 Conclusion

In this paper, we first demonstrate the main benefit of standardizing effect size estimates when reporting single fMRI studies. As standardized effect magnitudes have no unit, it becomes possible to compare the reported estimates over studies. In contrast, the BOLD percent signal change depends on the technique to calculate an average (i.e. baseline) signal which varies with software packages. We then discuss two main techniques to aggregate fMRI data over studies (i.e. a meta-analysis). The first one is the extension of the two-stage GLM approach (typically used in the analysis of single fMRI studies) with a third level. Ideally, the data is modelled using a mixed-effects approach (incorporating both within- and between-study variability). One benefit here is the usage of the original data. No transformation of the original data is needed before the meta-analysis. However, the analyst needs to be careful for unit mismatches in the model between studies in the database. This might occur when pre-processing of the individual studies is done using different software packages. An alternative technique avoids this problem by first transforming the reported study-level effect sizes to standardized effect sizes. These are then modelled again using a mixed-effects approach, incorporating both within- and between-study variability. Studies with small sample sizes (i.e. higher within-study variability) are down-weighted when estimating the population effect. Finally, we show that considerable between-study heterogeneity (which is not explained by moderator variables) can be expected in whole-brain fMRI meta-analyses.

We hope to further encourage data sharing practices so that complete fMRI images can be aggregated using appropriate methods.

## Conflict of interest

The authors declare no conflict of interest

## 5 Appendix

### 5.1 Distribution of a standardized mean effect

In this section, we provide the expectation and variance of a standardized mean effect. The derivations are obtained using Hedges (1981) which provides the expectation and variance of a standardized mean difference between an experimental and control group. The following can be used for estimating a standardized mean effect in one sample.

First consider a response variable *Y* of study *i* with *i* = 1, …, *k* and sample size *n*_*i*_. We assume 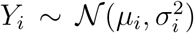. Furthermore, we are interested in estimating the mean effect 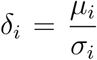 using 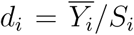, where *S*_*i*_ is the square root of the unbiased sample variance of study *i*. Note the following property of the sample variance, *S*^2^:

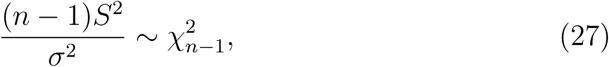

where *χ*^2^ is a Chi-squared random variable with *n* − 1 degrees of freedom (df).

Before we derive the expectation and variance of *d*, consider the expression of a non-central *t*-distributed random variable:

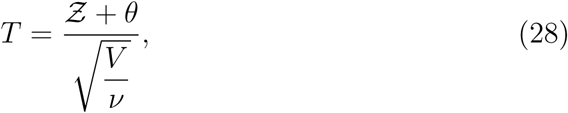

where 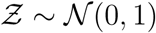, *V* is a Chi-squared distributed random variable with *n* − 1 degrees of freedom and *θ* is the non-centrality parameter.

Now we re-write the expression for *d* where we drop the subscript *i* for ease of notation.

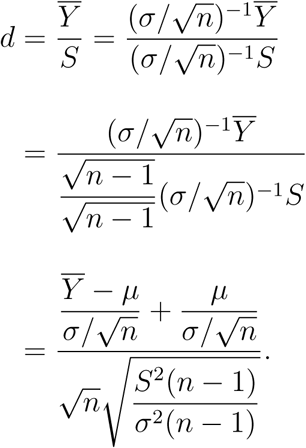

Denote 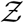 again as a standard normal distributed random variable. Furthermore using property (27), we write 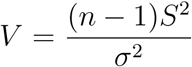 and *ν* = *n* − 1. Hence we get:

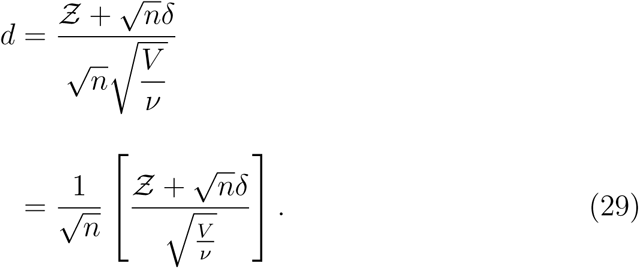

Using equation (28), it follows that *d* equals 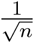 times a non-central *t*-distributed random variable with *n* − 1 degrees of freedom and 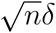 as the non-centrality parameter.

Using expression (29) and the first moment of the non-central *t*-distribution (Johnson and Welch, 1940), we have for *ν* > 1:

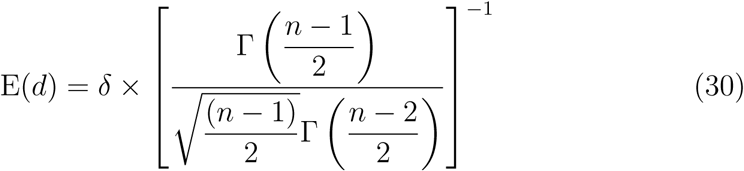

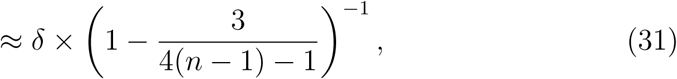

where Γ is the gamma function: Γ(*x*) = (*x* − 1)!

Furthermore, we define:

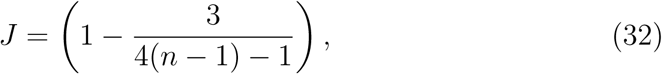

which is an approximation for 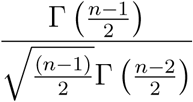 in equation (30) (Hedges, 1981).

When *n* ≥ 10, then 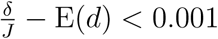.

For *ν* > 2, the second moment of the non-central *t*-distribution, gives:

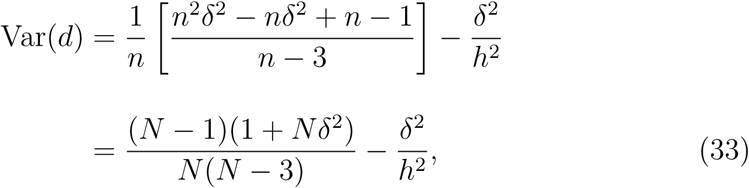

where

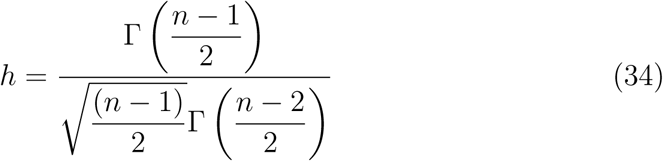

Note that it is again possible to replace *h* with *J*. Finally we plug in the estimate for *δ* in both equations (31) and (33).

As is clear from expression (31), 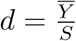 is a biased estimator for *δ*. For this reason, Hedges (1981) suggested to use:

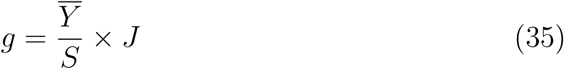

as an unbiased estimator. The variance then simply becomes:.

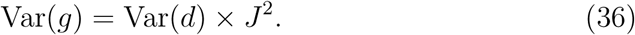

An alternative is the exact expression:

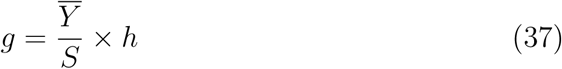

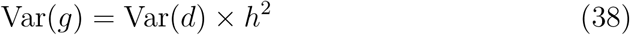

### 5.2 Empirical variance of estimator for *g*_*e*_

**Figure 7:**
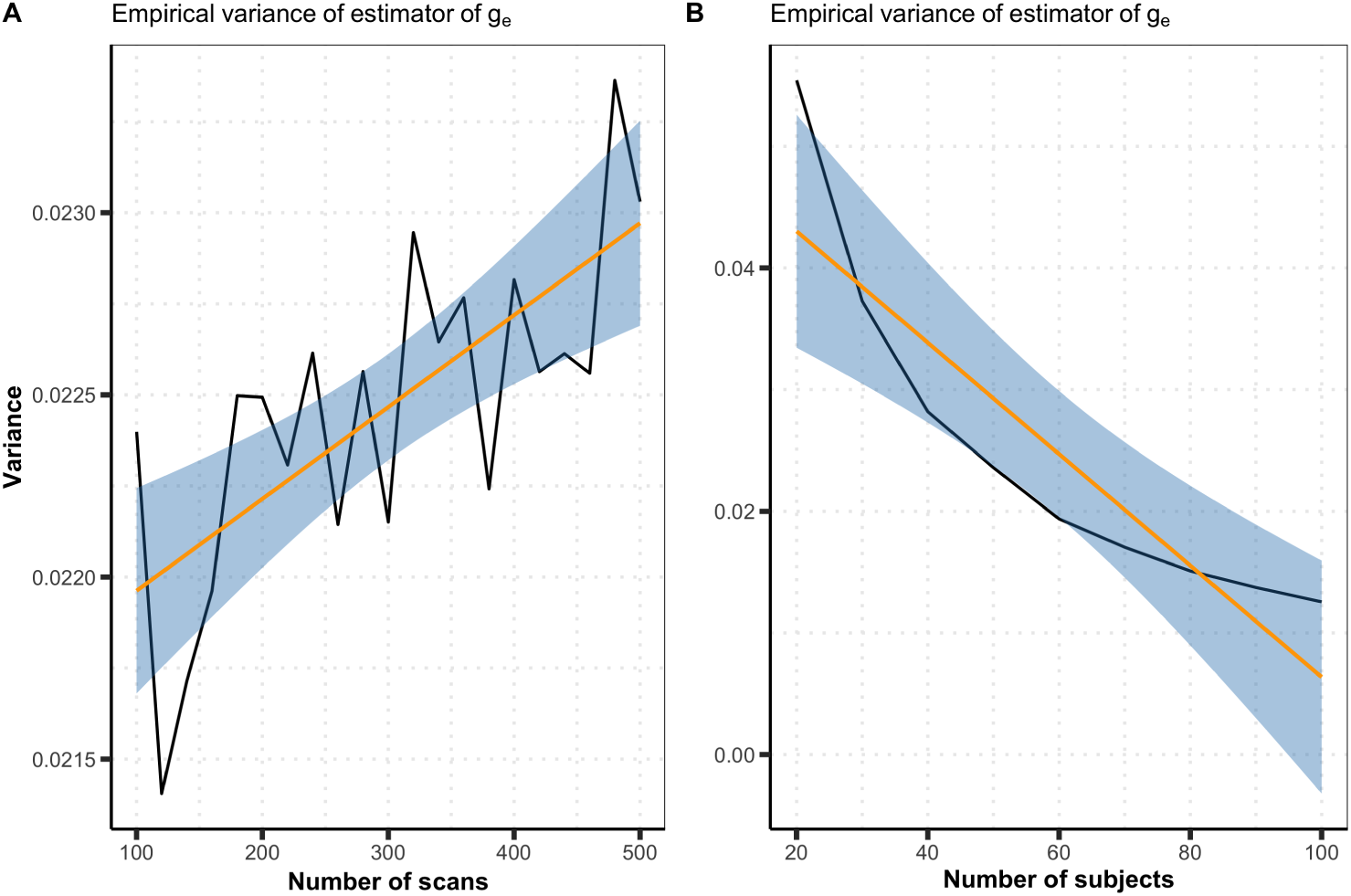
Empirical variances of the estimator for *g*_*e*_. Variance is calculated on the estimates of *g*_*e*_ obtained in the 1000 Monte-Carlo simulation runs. In panel **A**, we calculate the variance marginally over the number of subjects. In panel **B**, we calculate the variance marginally over the number of scans.

### 5.3 Unbiased standardized effect size estimator

**Figure 8:**
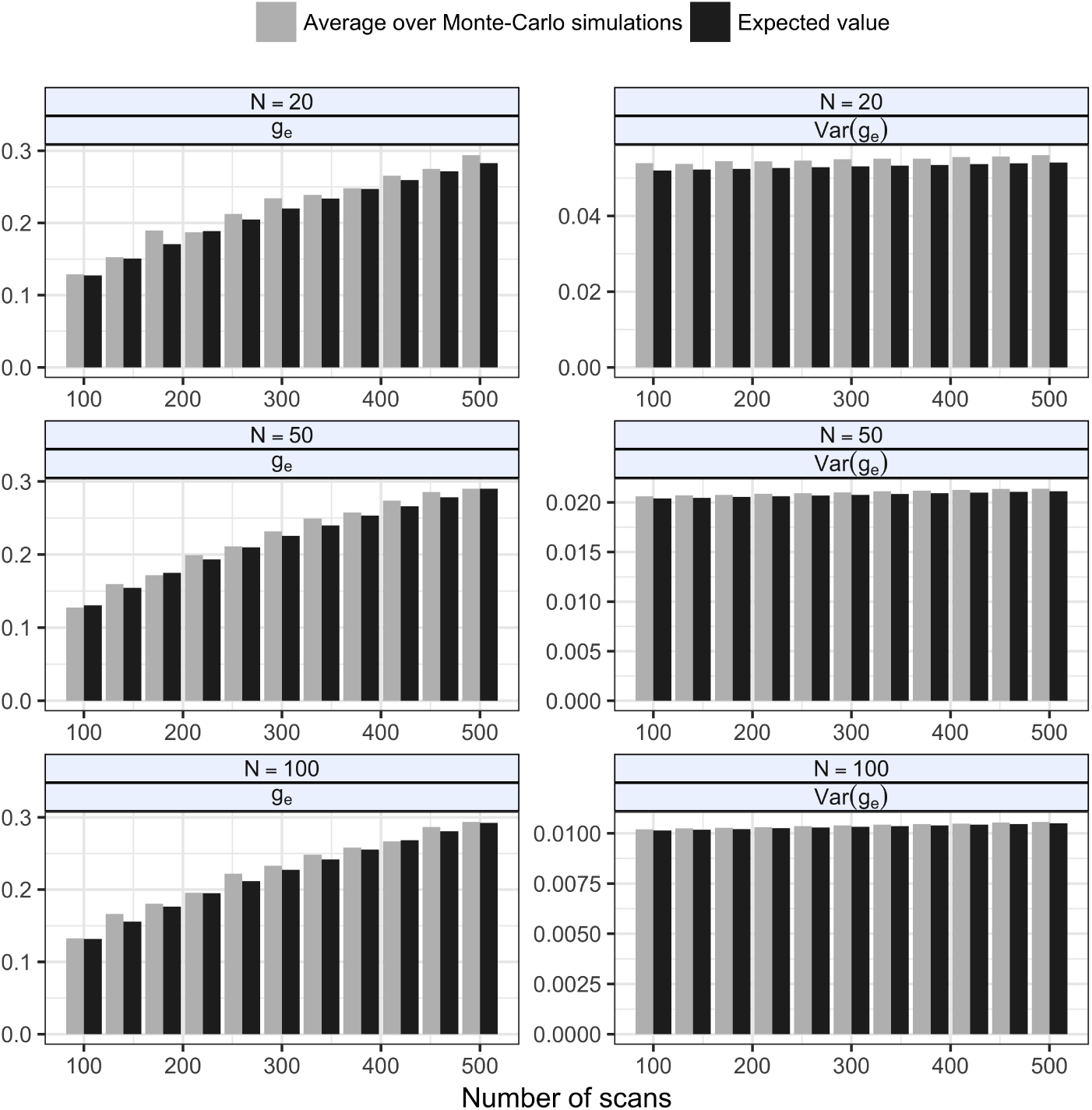
Empirical assessment of bias when estimating Hedges’ *g*_*e*_ and its variance. Results are obtained using 1000 Monte-Carlo simulations. Individual fMRI time series are generated with Gaussian noise and homogeneous within-subject variability. Group level model parameters are estimated using the OLS estimators. Both the average over all simulations and the expected values are plotted for different time series lengths and sample sizes.

### 5.4 Estimators for *τ* ^2^

To begin with, we have 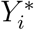, the (standardized) effect size from study *i* with *i* = 1,…, *k*. Next, 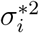 is the corresponding within-study sampling variance and a weight 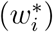 for each study is defined as 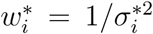. The weighted average, 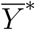, is defined as

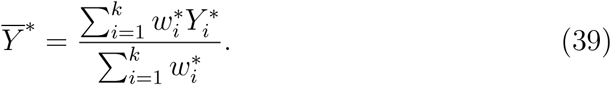

The first estimator for *τ* ^2^ is the DerSimonian and Laird estimator (DerSimonian and Laird, 1986):

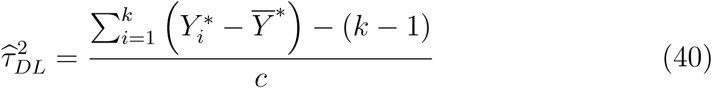

with

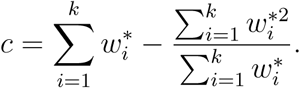

The second estimator for *τ* ^2^ is the Hedges estimator.

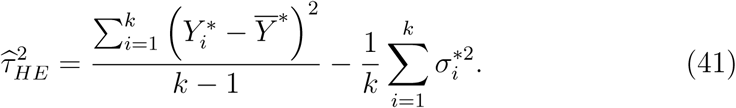

It has been shown that the Hedges estimator in equation (41) is unbiased, even when substituting unbiased estimates for the 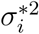 values (Viechtbauer, 2005).

The final estimator is based on the Restricted Maximum Likelihood approach (REML). In the approach presented here, the weights 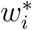 are first defined as 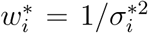, but after estimating 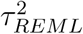 updated to 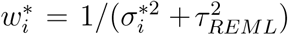. The estimator is given by

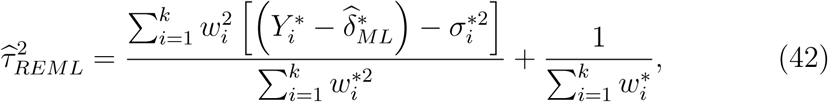

where 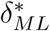 is the maximum likelihood estimator equal to expression (39). The estimator for 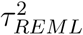 is obtained by iterating between 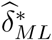 and 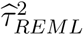 until convergence.

## Footnotes

1 http://neuropowertools.org

2 http://www.fil.ion.ucl.ac.uk/spm

3 https://fsl.fmrib.ox.ac.uk/fsl/fslwiki

4 https://blog.nisox.org/2012/07/31/spm-plot-units/

5 https://www.fil.ion.ucl.ac.uk/spm/software/spm12/

6 The code can be found at: https://github.com/NeuroStat/ESfMRI

7 http://www.prisma-statement.org/

8 http://neurosynth.org/

9 https://github.com/NeuroStat/NeuRRoStat

## Notes

https://github.com/NeuroStat/ESfMRI

